# Aberrant remodelling of astrocytic architecture in acute hepatic encephalopathy: complexity of oedematic atrophic astrocytes

**DOI:** 10.1101/2024.07.08.602491

**Authors:** Mariusz Popek, Marta Obara-Michlewska, Łukasz Szewczyk, Marcin Kołodziej, Alexei Verkhratsky, Jan Albrecht, Magdalena Zielińska

**Affiliations:** Department of Neurotoxicology, Mossakowski Medical Research Institute, Polish Academy of Sciences, Pawińskiego St. 5, 02-106 Warsaw, Poland; Laboratory of Molecular Neurobiology, Centre of New Technologies, The University of Warsaw, Banacha St. 2c, 02-097 Warsaw, Poland; Institute of Theory of Electrical Engineering, Measurement and Information Systems, Warsaw University of Technology, Koszykowa St. 75, 00-662 Warsaw, Poland; Faculty of Biology, Medicine and Health, The University of Manchester, Manchester, UK; Department of Neurosciences, University of the Basque Country UPV/EHU, 48940-Leioa, Spain & IKERBASQUE, Basque Foundation for Science, Bilbao, Spain; Research Centre on TCM-Rehabilitation and Neural Circuit, School of Acupuncture and Tuina/Health and Rehabilitation, Chengdu University of Traditional Chinese Medicine, Chengdu, China; Department of Forensic Analytical Toxicology, School of Forensic Medicine, China Medical University, Shenyang, China; Department of Stem Cell Biology, State Research Institute Centre for Innovative Medicine, LT-01102, Vilnius, Lithuania

**Keywords:** astrocyte, acute liver failure, aquaporin 4, ezrin, profilin-1, perisynaptic astrocytic processes, glutamate

## Abstract

Hepatic encephalopathy (HE) following acute liver failure (ALF) is a primary toxic astrocytopathy, although in-depth characterisation of underlying pathogenesis is far from complete. Among the multitude of astrocyte-specific proteins guiding brain functionality, plasmalemma-cytoskeletal linker ezrin, actin-binding protein profilin-1, and water channel aquaporin 4 (AQP4) contribute to astrocytic morphological plasticity through regulation of cell shape, volume, complexity of primary and terminal processes, and positioning astrocytes against other CNS constituents. Changes in these proteins might contribute to the brain oedema and astrocytic morphological remodelling in the HE. Using transmission electron microscopy, confocal fluorescent microscopy, and 3D reconstruction, we found complex morphological alterations of cortical astrocytes in mice with azoxymethane-induced ALF. Astrocytic primary branches demonstrated hypertrophy, whereas terminal leaflets showed atrophy quantified by the reduced area occupied by astrocytes, decreased number and the length of leaflets, decreased leaflets volume fraction, and altered astrocyte-to-neurone landscape. These morphological changes correlat with decreased expression of AQP4, phosphorylated leaflet-associated ezrin, and the actin dynamics regulator, profilin 1, suggesting the contribution of these proteins to astrocytic pathological remodelling. Pathological changes in astrocytes develop in parallel, and are likely causally linked to, the HE-linked neurological decline, manifested by a reduction in electroencephalography power and by excessive glutamate in the brain microdialysates.

Graphical abstract:
Hepatic encephalopathy is associated with astrocyte remodelling manifested by swelling of the soma and primary branches together with atrophy of distal branches and leaflets; the latter retract from synapses thus affecting neurotransmission and contribute to the reduced neuronal activity. Astrocyte remodelling was linked to (and probably instigated by) a decrease of plasmalemma-cytoskeleton linker phosphorylated ezrin (Phos-ezrin), actin modulator profilin-1 (PFN1) and water channel aquaporin 4 (AQP4).

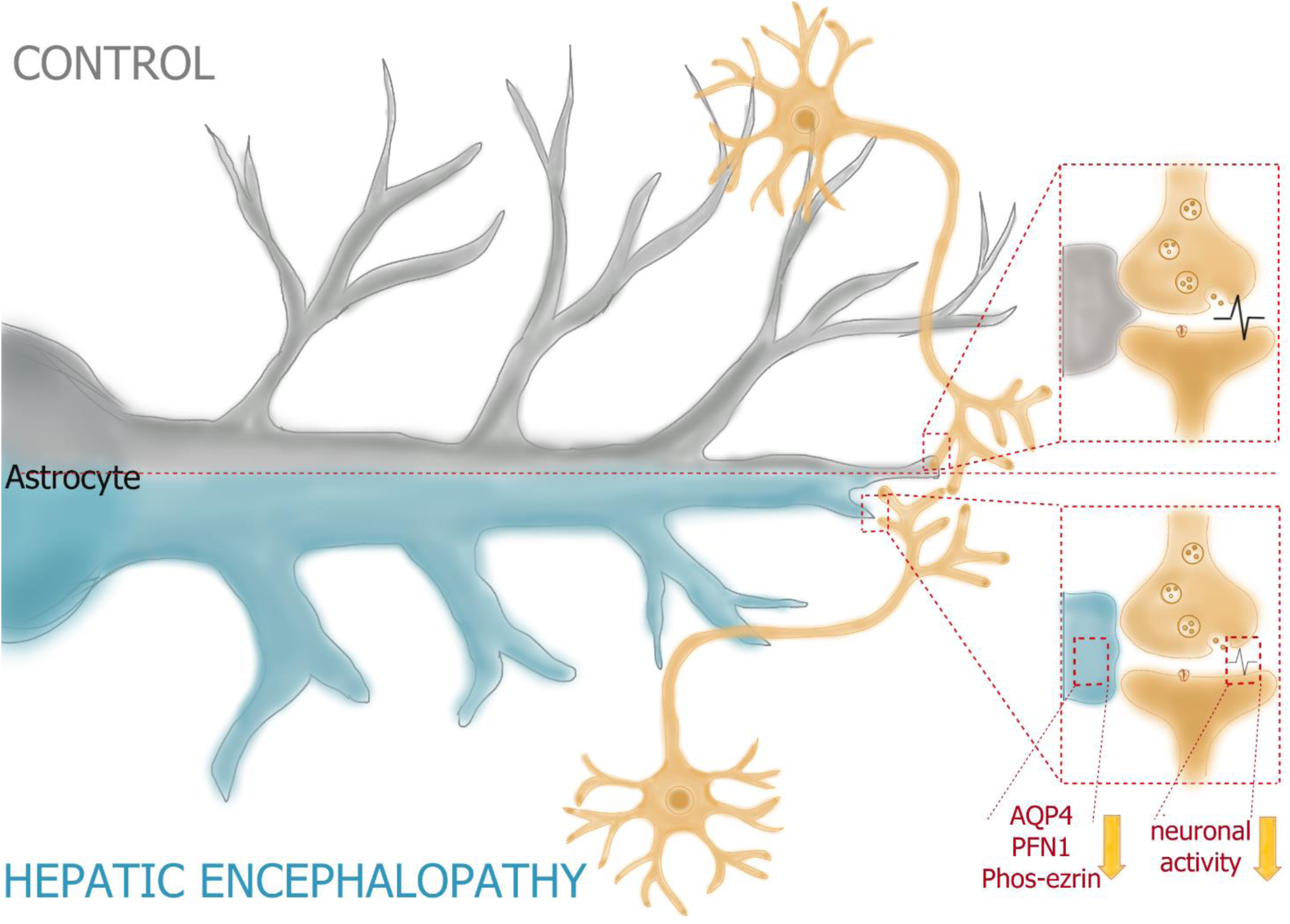

## Introduction

Hepatic encephalopathy (HE), the primary astrocytopathy (Butterworth, 2022; Verkhratsky et al., 2023), is a complex, multi-symptomatic neurological syndrome caused by a systemic increase in ammonium (Aldridge et al., 2015) usually linked to an acute or chronic liver failure (ALF and CLF respectively). Ammonium circulating in excess enters the brain, where it is detoxified and converted to glutamine by astrocyte resident glutamine synthetase (Brusilow et al., 2010; Cooper et al., 1979). Pathological increase of glutamine in the brain leads to a multifaceted homeostatic imbalance (Zielińska et al., 2022), which is clinically manifested by delirium, hallucinations, acute psychosis, and, in a terminal phase, coma leading to death (Weissenborn, 2019). In HE astrocytes undergo multiple morphological changes (Albrecht & Jones, 1999; Gelpi et al., 2020) ranging from reactive astrogliosis, swelling (that contributes to brain oedema), and the emergence of disease-specific aberrant astrocytes known as Alzheimer type II astrocytes, the term introduced in 1942 (Waggoner & Malamud, 1942).

Acute liver failure (ALF) is employed as an experimental model of HE. Animals subjected to ALF demonstrate prominent neurological decline and synaptic impairments (Hamdani et al., 2021; Popek et al., 2018), associated with brain oedema as well as the abnormal flow of the cerebrospinal fluid (Obara-Michlewska et al., 2018). Experimental observations revealed that ionic composition and acid-base balance of the interstitial fluid remain well controlled (Obara-Michlewska et al., 2018), a feature possibly contributing to the partial brain function recovery observed in human patients (Anand & Singh, 2019; Ochoa-Sanchez et al., 2021).

Protoplasmic astrocytes are characterised by a complex morphology defined by an extensive arborisation comprised of primary processes known as branches and distal exceedingly thin processes defined as leaflets (Khakh & Sofroniew, 2015; Semyanov & Verkhratsky, 2021). Leaflets, often associated with synapses, demonstrate remarkable morphological plasticity relevant to synaptic function (Henneberger et al., 2020). Morphological plasticity of astrocytic processes is regulated by the leaflet-resident ezrin, a linker protein, connecting plasmalemma with the actin cytoskeleton (Derouiche & Geiger, 2019; Lavialle et al., 2011). Astrocytic arborisation may also be regulated by profilins linked to actin dynamics. In particular, profilin 1 (PFN1) was shown to regulate astrocyte stellation (Schweinhuber et al., 2015) and Ca^2+^-induced extension of leaflets in cultured astrocytes (Molotkov et al., 2013). Another astrocyte-specific protein associated with the regulation of astrocyte volume and morphology is the water channel aquaporin 4 (AQP4) (Tang & Yang, 2016), which, in the healthy brain, is primarily localised to astrocytic endfeet plastering blood vessels (Nagelhus & Ottersen, 2013). Two classical isoforms, AQP4a (M1) and AQP4c (M23) are alternatively spliced transcripts of the *AQP4* gene that participate in the cell volume regulation of astrocytes (Jorgačevski et al., 2020). The M23 isoform is also involved in the remodelling of the astrocytic processes, especially of those in the vicinity of the glutamatergic synapses (Ciappelloni et al., 2019).

An in-depth analysis of astrocytic branches and leaflets phenotypes and synaptic landscape in the HE, however, has not been performed. Here we present data indicating complex changes in morphology of protoplasmic astrocytes in the HE induced by ALF. On one hand, astrocytic branches and soma increase their presence, on the other we found a prominent atrophy of the perisynaptic leaflets leading to decreased synaptic astrocytic coverage and altered astrocyte-synaptic landscape. These morphological aberrations were paralleled by substantial changes in the astrocytic content of phosphorylated ezrin, PFN1, and AQP4. Morphology and protein changes correlate with overall inhibition of the brain activity documented by EEG, reflecting loss of astrocytic homeostatic function and pathological astrocyte-neuron communications that may be directly associated with HE neuropathology.

## Material and Methods

### Animal models

Experiments were performed on 10–12 weeks male C57Bl mice from an animal colony of the Mossakowski Medical Research Institute, Polish Academy of Sciences, and on Aldh1l1_Cre_ERT2 :TdTomatoWT/fl mouse line Control TdT+] (form the colony of Centre of New Technology). All experimental procedures conformed EC Directive 86/609/EEC, with approval and under surveillance of the IV^th^ Local Ethical Committee in Warsaw, Poland.

### Azoxymethane model of ALF in mice

ALF in mice was induced by a single intraperitoneal (i.p.) administration of hepatotoxin azoxymethane (AOM), (Sigma-Aldrich, A5486, Poznań, Poland) at 100 mg/kg body mass. The body temperature of animals was maintained with the heating pad (37°C), whereas the hypoglycaemia and dehydration were prevented with two (8 and 16 hours post-AOM) i.p. injections of 500 μl of 5% glucose and 0.18% saline solution (Popek et al., 2018).

### Hematoxylin and eozin liver staining

At the selected time points post AOM injections (4, 12, 18, 24h) mice livers were exposed and examined with light microscopy. Subsequently, the liver was subjected to hematoxylin and eozin (H&E) staining. Briefly, tissue was frozen in dry-iced 99.5% acetone (Sigma-Aldrich, Poznań, Poland) at −80°C. Livers were cut into 10 µm thick sections (Leica CM1860 cryostat), and washed with an ethanol gradient (99%-70%) and water. Sections were stained with hematoxylin (Sigma Aldrich, St. Louis, MO, USA) solution and rinsed under running water for 10 min, followed with 4 min incubation with eosin (Sigma Aldrich, St. Louis, MO, USA) solution, and washed with an inverted ethanol gradient. The sections were mounted in DPX medium (Sigma-Aldrich, 06522, Poznań, Poland) and analysed using an Olympus IX71 Inverted Fluorescence Motorised Microscope (Olympus Corporation, Tokyo, Japan).

### Serum biochemical analysis

Concentration of serum glucose, ammonium, alanine aminotransferase (ALT), total protein, cholesterol, and alkaline phosphatase (ALP) in Control, 4, 12, 18, and 24 hours post AOM injection mice were analysed using IDEXX Catalyst One (IDEXX Laboratories, Inc., Westbrook, Maine, U.S.A.). Briefly, blood was immediately collected to heparin tube and according to IDEXX protocols, Chem 15 CLIP were used. Separately, maintaining a 2-hour interval between other sample measurements, the ammonia concentration was measured immediately after plasma collection, using an Ammonia individual slide (IDEXX Laboratories, Inc., Westbrook, Maine, U.S.A.).

### Transmission electron microscopy

Anaesthetised mice (2% isoflurane) were perfused through the ascending aorta with 2% paraformaldehyde and 2.5 % glutaraldehyde in 0.1 M cacodylate buffer, pH 7.4. The frontal cortex of control and HE mice (24 h post-AOM injection) was fixed in the mentioned solution for 24 h (at 4°C) and placed in a mixture of 1% OsO4 and 0.8% K_4_[Fe(CN)_6_], as previously described (Popek et al., 2018). Astrocytic content in the neuropil of the frontal cortex was estimated from 15 randomly selected sections obtained from each of the three control mice and 15 sections from four AOM mice. Sections with clearly visible fragments of cell bodies were excluded, but the remaining astrocytic processes were not divided morphologically, apart from the perivascular ones. Following parameters were quantified: astrocytic area content on each section, the average number of astrocytic leaflet apposition to the synaptic cleft, the average distance between a proximal end of the active zone (AZ) and the nearest astrocytic leaflet, the average distance between AZ to nearest AZ and composition of neuropil along AZ to nearest AZ paths. For image analysis, ImageJ processing software was used. Representative sizes and types of 268 cortical synaptic complexes from control and 268 from HE sections were analysed.

### Real-time PCR

In the homogenates of mouse cerebral cortex, at indicated time points after AOM administration, the mRNA level of AQP4 and its isoforms, M23 and M21, was assessed with real-time PCR in Applied Biosystems 7500 Fast Real-Time PCR System (Applied Biosystems, Foster City, CA, USA). Total RNA was extracted according to the phenol-chloroform method (RNA Extracol, Eurx, Gdańsk, Poland) (Chomczynski & Sacchi, 1987)). RNA samples (1 μg) were reversely transcribed (High-Capacity cDNA Reverse Transcription Kit; Thermo Fisher Scientific, Waltham, MA, USA) and real-time PCR analysis was performed using 1μl of cDNA in a reaction of 10 μl. Fast SG/ROX qPCR Master Mix and SYBR Green Fast SG qPCR Master Mix (Eurx, Gdańsk, Poland) were used for the analysis of total AQP4 transcript, and M23 and M1 isoforms, respectively. TaqMan assay IDs were Mm00802131_m1 for Aqp4 and Mm00607939_s1 for β-actin (Thermo Fisher Scientific, Waltham, MA, USA), whereas primer sequences for M1 and M23 isoforms were synthesised according to (Fallier-Becker et al., 2016) (Fallier-Becker et al., 2016). Data are averaged triplicates of mRNA transcript levels, normalised to β-actin expression according to the 2-ΔΔCt method (Livak & Schmittgen, 2001).

### Western blot

Tissue samples were collected in RIPA buffer (Thermo Fisher Scientific, Waltham, MA, USA), supplemented with 0.05% CHAPS detergent and protease/phosphatase inhibitors cocktail (1:100) (Merck, Darmstadt, Germany), homogenised, sonicated and centrifuged (10 min, 10 000 g, 4 °C). Protein content in the supernatant was quantified using Pierce™ BCA Protein Assay Kit (Thermo Fisher Scientific, Waltham, MA, USA; Cat No.: 23225). Thirty micrograms of protein were run on a 10 % SDS-polyacrylamide gel (Thermo Fisher Scientific, Waltham, MA, USA) and then transferred onto a PVDF membrane with semi-dry Trans-Blot Turbo System (Bio-Rad Laboratories, CA, USA). Blots were blocked in 5% bovine serum albumin (BSA) (BioShop; Lab Empire S.C.) and incubated in 2.5 % BSA, overnight, 4 °C, with primary antibody: Aqp4 (1:1000; PA5-85767, Thermo Fisher Scientific, Waltham, MA, USA), PFN1 (1:1000; P7624, Merck, Darmstadt, Germany), ezrin (1:1000; SAB4200806, Merck, Darmstadt, Germany). Protein bands were detected with host-specific secondary antibodies and developed with a chemiluminescent substrate (GE Healthcare Amersham, Piscataway, NJ, USA). After stripping, the blots were incubated with HRP-conjugated anti-GAPDH antibody (1:8000; HRP-60004, Proteintech, Manchester, UK) and developed. The chemiluminescent signal acquisition and densitometric analysis were conducted using the G-Box system (SynGene) and GeneTools software (SynGene), respectively.

### Brain preparation and immunohistochemistry

The brains of C57Bl mice, divided into control and HE (4, 12, 18, 24 h AOM post-injection) groups, were quickly removed, immediately frozen in powdered dry ice, and stored at −80°C. 20 µm thick frontal cortex sections (Leica CM1860 cryostat) were proceeded according to standard protocol. Briefly, after 20 min of 4% PFA fixation, 1 h blocking with normal goat serum (10%, diluted with 0.1% Triton X-100), overnight incubation with Phospho-Ezrin antibody (Cell Signaling Technology, 3726T, Massachusetts, USA) 1 h incubation with goat anti-rabbit IgG Alexa Fluor 546 secondary antibody (1:500, Life Technologies, Carlsbad, California, USA) were performed. For astrocytes labelling, GFAP primary antibody (1:500, sc-33673, Santa Cruz Biotechnology, Texas, USA) and goat anti-mouse IgG Alexa Fluor 488 secondary antibody (1:500, Life Technologies) were used. For cell nuclei staining 5 μM Hoechst (1:500, Life Technologies, 33258) was used. Procedures for negative controls were carried out in the same manner except that the primary antibodies were omitted. Z-Stack photos were conducted using a confocal laser scanning microscope (LSM 780, Carl Zeiss Meditec, Germany).

For fluorescence intensity analysis of astrocyte processes and quantification of astrocytes single branches and neuronal volume, astrocyte TdT+ mice line was used. Control TdT+ and 24h post AOM injection HE mice were transcardially perfused with 0.1 M PBS, followed by 4% PFA (Sigma Aldrich, 158127) in PBS. Brains were dissected, incubated overnight in 4% PFA, and saturated with 30% sucrose (Merck Life Science, cat. no. 1076515000) in PBS at 4 °C for the next 24 hours. Next, the brains were transferred into O.C.T. (Sakura Finetek Europe B.V., 4583) and frozen in −30°C isopentane (VWR, 24872.298). Frontal cortex 30 μm sections (Leica CM1860 cryostat) were prepared as free-floating sections in anti-freeze solution (30% sucrose (Merck Life Science, 84100); 30% glycerol (VWR, 443320113); 0.1 M PBS, pH 7.4). After PBST washing, antigen retrieval was conducted using 10 mM sodium citrate buffer (Bioshop Life Science Products, CIT001.1) with 0,05% Tween 20 (Sigma Aldrich, P7949). Next, sections were blocked 1h in 10% donkey serum in PBST and incubated overnight with primary antibodies for anti-GFAP (1:250, Merck Life Science, Hpa056030) and anti-MAP2 (1:200, Sigma-Aldrich, M3696). Further, the slices were washed and incubated with goat anti-rabbit IgG Alexa Fluor 488 secondary antibody (1:500, Life Technologies, Carlsbad, California, USA). Finally, slices were mounted on glass slides with Fluoromount G. Z-Stack photos were conducted using an Axio Imager Z2 LSM 700 Zeiss confocal microscope.

### Immunofluorescence analysis

Z-Stack images were processed using the ZEN v. 2008 (Carl Zeiss Meditec, Germany). The fluorescence intensity of Phos-ezrin was analysed from the mean grey value of Maximum Intensity Z-projected images using Image J software and normalised to analysed the astrocytes area.

### Astrocytic processes distribution analysis

The ring shaped ROIs with 5 µm decrement were applied to Maximum Intensity Z-projected images of astrocytes. The fluorescence intensity of one astrocyte was analysed in each of the 6 ROIs and normalised to the fluorescence intensity of the individual, not-saturated soma. The ROI fluorescence intensity in distance between 15-20µm from the soma was analysed as characteristic of fine processes and considered as a volume fraction (VF) parameter. Analysis was based on the mean grey value of Z-Stack images using Image J software.

### Sholl analysis of astrocytes

Astrocytes Z-stack images were traced and skeletonised using SNT plug to Fiji. Sholl analysis was performed from soma centre every 2µm and number of intersections was counted. Based on skeletonised model, number of junctions (treated as points from which at least 2 branches originate) and length of the longest branch were calculated.

### Astrocytes and neurons volume quantification

The volume of astrocyte single branch and neurones in the same ROI was measured using the MeasurementPro, Imaris 8.4.2 software (Bitplane AG). Briefly, three-dimensional surface rendering of individual astrocytes was generated, and within the area occupied by a single astrocytic branch, MAP2-staining-based neuronal surfaces were constructed. Renders were created based on the pixel gradient intensity algorithm, using 488 nm and 546 nm channels. Renders were threshold to ensure all processes of astrocytes were properly reconstructed. Pictures were blinded before analysis.

### EEG recording of the freely moving mice

Mice were anaesthetised with ketamine (75 mg/kg, Vetoquinol) and dexmedetomidine (0.5 mg/kg, Orion Pharma) for stereotactic implantation of electrodes. The 2EEG/1EMG headmounts (EEG/EMG/Plus Mouse Headmount, Pinnacle Technology, Lawrence, KS) were affixed to the exposed skull with stainless steel screws acting as EEG electrodes. The screws were positioned to be resting on the meninges. Two parietal screws were used for EEG recording and one frontal screw for reference and grounding. EMG electrodes were implanted below the neck muscles. The headmount was secured with dental acrylic cement and the skin incision was sutured below and above the headmount. Following the surgery, mice were woken up from anaesthesia with atipamezole hydrochloride (1 mg/kg, Orion Pharma) and treated with ketoprofen (5 mg/kg, Sandoz) once a day for up to 3 days.

EEG recordings were conducted in a transparent, round cage with free access to food and water, to which mice were acclimated for one day. For EEG recording, the mice were connected to the data acquisition system (8401 Data Conditioning & Acquisition System, Pinnacle Technology), with the attachment plug tethered to the swivel at the balanced arm that, together with the revolving of the cage, allowed animals for moving freely during the EEG recording. The 3 – 4 hours recording was acquired, using Sirenia software (Pinnacle Technology), before AOM administration. The recording was carried on within 24 hours of ALF development.

For each EEG data set signal power was calculated in particular frequencies using Butterworth filters (Kadam et al., 2017): total EEG band (0–100 Hz), delta (1–4 Hz), theta (4– 8 Hz), alpha (8–13 Hz), beta (13–30 Hz), gamma (30–100 Hz), high-frequency oscillations (HFO) (>80 Hz). Thirty-minute epochs, baseline, and overlapping the 4, 12, 18, and 24 h time points after AOM administration were analysed by Matlab program. The power values were normalised and presented as relative to the baseline.

### Microdialysis of freely moving mice

A microdialysis cannula was implanted into the frontal cortex (coordinates: AP +2.0, ML - 0.8, DV −1.5) under isoflurane (Aerrane, Baxter) anaesthesia, and mouse recovery time was set as two days. After this time, CMA7 microdialysis probe (2mm, Harvard Apparatus, Massachusetts, United States) with aCSF (composition in mM: NaCl (130), KCl (5), CaCl_2_ (2.5), MgSO_4_ (1.3), KH_2_PO_4_ (1.25), NaHCO_3_ (26), and D-glucose (10)) flowing through at a rate of 2.5ul/min, was implanted to control or AOM-induced mice. Microdialysates from freely moving mice brains were collected every 40 minutes (100 µL) and immediately frozen at −80°C.

### High-performance liquid chromatography

Glutamate extracellular concentration was measured in dialysates using HPLC with fluorescence detection after derivatisation in a timed reaction with o-phthalaldehyde with mercaptoethanol, as described earlier (Hilgier et al., 1999). Samples (50μL) were then injected into a 5μm Bio-Sil C18 Hl column (250 × 4.6 mm, BIO-RAD), with a mobile phase of 0.075 M KH_2_PO_4_ solution containing 10% v/v methanol, pH 6.2 (solvent A), and methanol (solvent B). Concentrations were averaged from the 2 fractions closest to the specified time point.

### Statistical analysis

All experiments were carried out with replicates depending on the experiment type. When two population groups of responses were examined, depending on the results obtained, the Student’s t-test or the Mann-Whitney U test was applied (see legends to figures). Statistical significance was determined by one-way analysis of variance (one-way ANOVA) followed by Dunnett’s post hoc comparisons. Error bars represent the SD, which is specifically indicated, *P < .05, **P < .01, and ***P < .001. All statistical analyses were performed using GraphPad Prism 7 (GraphPad Software, Inc., USA).

## Results

### General pathology of HE mice

#### Biochemical and morphological characteristics of ALF mice

Progression of the ALF in mice was monitored at 4, 12, 18, and 24 hours after AOM injection. Gradual deterioration of biochemical parameters was measured in blood serum (Fig.1B), confirming the evolution of liver failure and HE symptoms. Histological examination of mouse livers exposed irregular and speckled liver surface observed as early as 4h, which gradually progressed through the later time points. Fine, bluish-red pattern of bleeding was observed at 12 h post-AOM; spotty surfaces depicting residual cirrhotic nodules were detected from 18 h post-AOM, and extensive necrotic spots were obvious at 24 h (Fig. 1A). Histopathological assessment with hematoxylin-eosin staining revealed tissue degeneration from 4 h, hepatocyte apoptosis, and partial hyperaemia at 12 h, followed by necrosis of hepatocytes, the fibre mesh scaffold collapse and inflammatory invasion at 18 h with damage furher increasing at 24 h (Fig. 1A).

**Figure 1.**
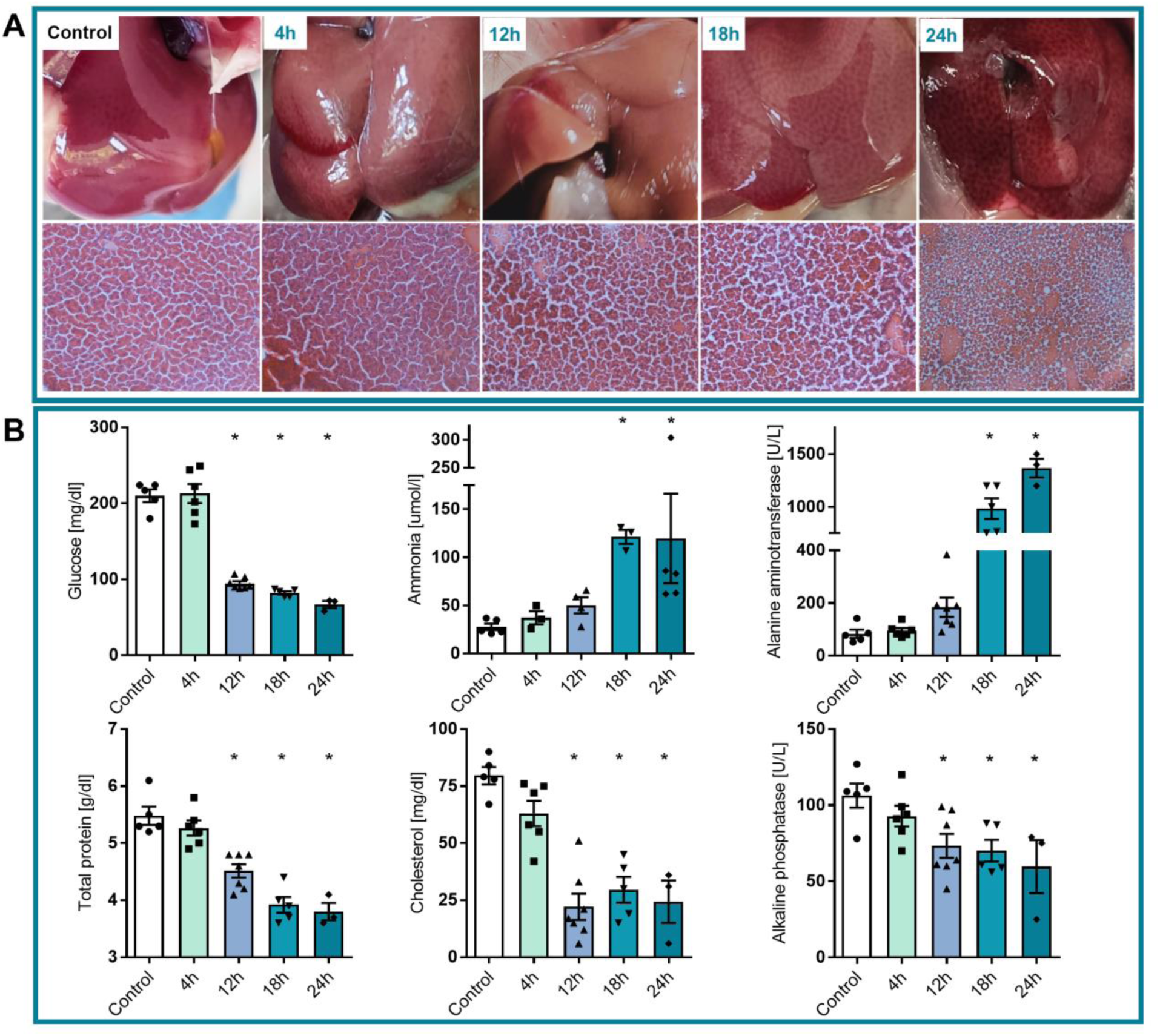
Validation of ALF mice model. A: Morphological and histopathological assessment of liver failure progression at 4, 12, 18, and 24 h after azoxymethane (AOM) injection. B: Serum levels of glucose, ammonium, ALT activity, total protein, cholesterol, and ALP activity at 4, 12, 18, and 24 h after AOM injection. n = 4-7, *p<0.05 ANOVA with Dunnett’s post hoc test.

Hyperammonemia (from 28 ± 7.4 μM in control to 120 ± 12.7 µM at 18 h and 24 h) and increased alanine aminotransferase (ALT) activity (from 84 ± 35 U/L in control to 984 ± 220 U/L at 18 h and 1366 ± 153 U/L at 24 h) were evident at 18 h after AOM injection (Fig. 1B). The serum glucose level decreased at 12 h (from 210 ± 18 mg/dl to 94.7 ± 7.5 mg/dl) after AOM administration (Fig.1B). Similarly, we observed decreased total protein (from 5.5 ± 0.4 to 4.5 ± 0.3 g/dl at 12 h and to 3.8 ± 0.3 g/dl at 24h), cholesterol (from 80 ± 8 mg/dl to 22 ± 15 mg/dl at 12 h and to 24 ± 16 mg/dl at 24 h), and alkaline phosphatase (ALP) activity (from 106 ± 18 U/L to 73 ± 21 U/L at 12 h and to 60 ± 30 U/L at 24 h) (Fig.1B).

### Electron microscope imaging of the HE mice brain cortex

To test whether ALF-induced HE modified astrocyte-synaptic landscape, we quantified cell ultrastructural morphometry. Astrocytic processes were identified by their irregular shape, the presence of glycogen granules and bundles of intermediate filaments in a comparatively clear cytoplasm (Fig. 2A). The astrocyte area in the section was determined by outlining and computing the area of all astrocytic processes and dividing by the total area of each section. Astrocytic endfeet in direct contact with blood vessels were counted separately from parenchymal processes classified as branches and leaflets. Sections with astrocytic cell bodies were not included in the analysis (1 section for the control group and 2 for HE). The area of astrocyte in relation to the area of section was counted; each cell body (either astrocyte with a visible nucleus or a neurone) occupying more than 20-25% of entire section introduces deviation, therefore the analysis of such section was subjectively excluded. In the control group, astrocytic endfeet occupied 2.85 ± 1.5% and astrocytic branches and leaflets 2.6 ± 1.2% of the total area while in HE mice they occupied, respectively, 6.3 ± 3% and 4.3 ± 1.7% (Fig. 2B). HE, therefore, is associated with a significant increase in astrocyte occupying territory, irrespectively from separate quantification of astrocytic endfeet or branches with leaflets.

**Figure 2.**
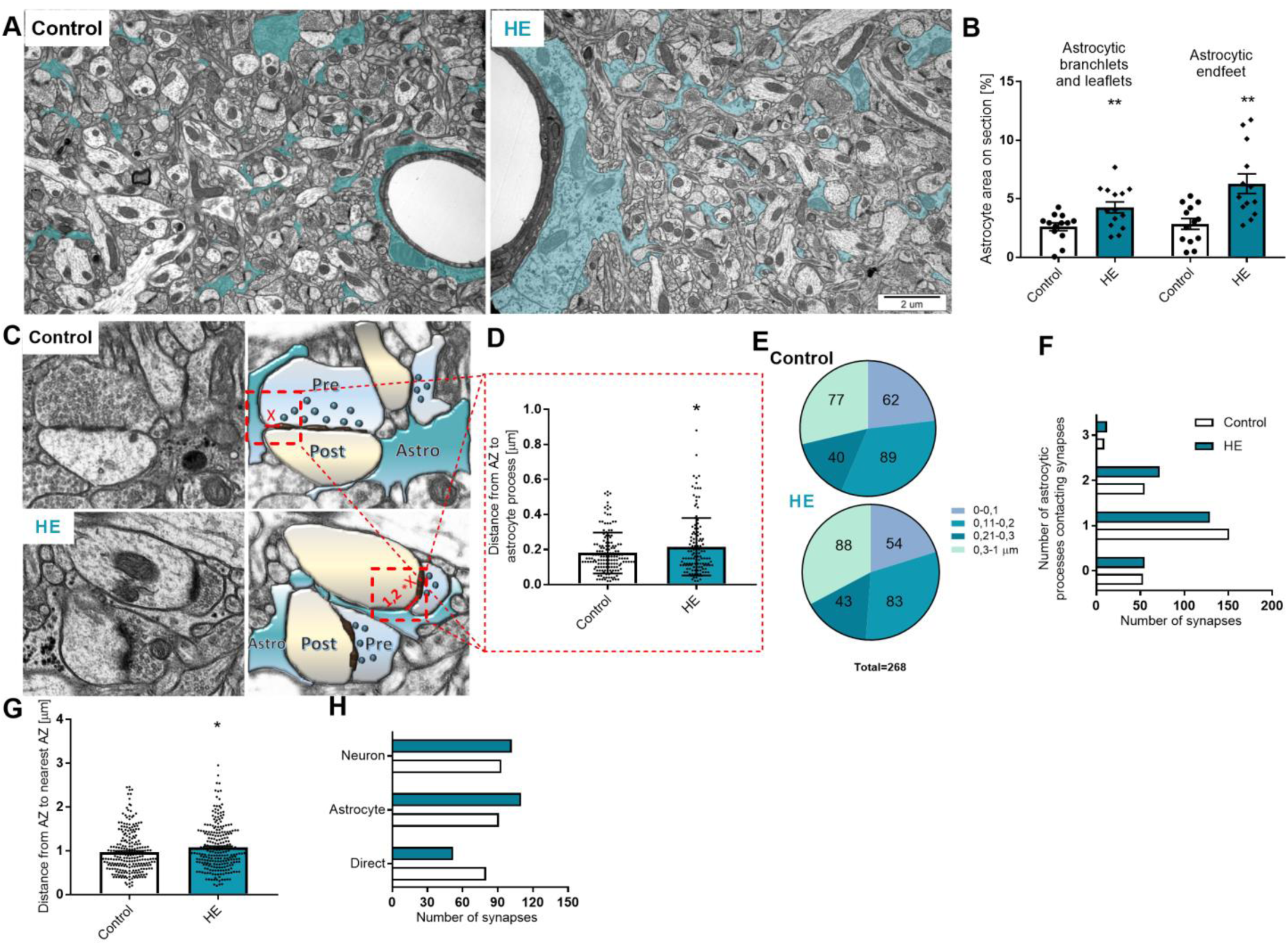
Ultrastructural pathohistology of HE astrocytes. A: Astrocytic profiles (navy blue) in a single thin section from control and HE (AOM 24 h) mice. B: Area of astrocytic processes referred to the whole analysed section from control and HE mice, n = 3 mice for control and 4 mice for HE, 13 sections for control and 14 sections for HE,* p < 0,05, t-Test. C: representative morphological interactions between astrocytic processes (Astro, navy blue) and synapses (presynaptic terminal – Pre, blue and postsynaptic bouton – Post, yellow) in control and HE mice; x denotes the distance between the active zone (AZ) edge and the nearest astrocytic membrane. D: Distance from AZ edge to the nearest astrocytic process, analysed in synapses which contact single astrocytic process (149 for control; and 129 for HE), n = 3 – 4 mice, *p < 0,05, t-test. E: Number of synapses in which the distance between AZ edge and astrocyte process was in the range between 0-100, 101-200, 201-300, and 300-1000 nm. F: Distribution of the number of astrocytic processes contacting synapses; 268 synapses from control and 268 synapses from HE mice. G: Distance from AZ edge to the nearest synapse AZ edge, 268 synapses were analysed from both groups, n=3 - 4 mice, *p < 0,05, t-test. H: Comparison between analysed synapses; (Directly – the nearest synapses are in direct contact, Astro – between the nearest synapses astrocytic processes is present, Neuro – neuronal component present between the nearest synapses), 268 synapses from n=3 - 4 mice were analysed.

Excluding ‘no contact synapses’ the distance between the nearest astrocyte process to the synaptic cleft (the distance to the proximal end of the active zone (AZ)) increased by ∼20% in HE group, from 0.18 ± 0.11 to 0.22 ± 0.16 μm (based on 151 and 129 synapses, respectively) (Fig. 2D). The same analysis binned to < 0.10 µm, 0.10 - 0.20 µm, 0.20 - 0.30 µm and > 0.30 µm indicates decrease in the number of astrocytic leaflets within the 0 - 0.10 and 0.10 - 0.20 µm distance from synapse and slight increase at > 0.3 µm distance range in HE compared to controls (Fig. 2E). The number of synapses with no contact with astrocyte and synapses contacted by 1, 2, or 3 astrocytic leaflets did not differ between groups (Fig. 2F). Distance between the nearest synapses measured as AZ edge to nearest AZ edge path length was 0.96 ± 0.47 μm in the control group and 1.07 ± 0.51 µm in the HE group (Fig. 2G).

The synaptic landscape was also qualitatively compared between control and HE animals. The comparison between analysed synapses indicates that 80 synapses in control and 52 in HE were in direct contact, whereas 91 nearest synapses from control and 110 synapses from HE were contacting with astrocytic processes, in turn, between 93 nearest synapses from control and 102 synapses from HE, neuronal component was observed in synaptic vicinity. (Fig. 2H).

### Changes in expression of aquaporin 4, profilin 1, ezrin, and phospho-ezrin in HE

We measured the protein level of AQP4, ezrin, and PFN1 and analysed fluorescence intensity of phosphorylated ezrin; (n = 11 and 3 in control and AOM-injected, respectively; n = 11 and n = 4 in control and AOM-injected, respectively; n = 7 and 3 in control and AOM-injected, respectively; n = 7 and n = 5 in control and AOM-injected, respectively). Protein level of AQP4, detected with an antibody specific against the C-terminus of both M1 and M23 isoforms, decreased by 42 % (from 107.6% ± 9.69 in control, to 62.55% ± 14.24) at 4 h (n = 7 and 3 in control and AOM-injected, respectively) and by 49% (to 58.98% ± 4.52) at 24 h post-AOM injection (n = 7 and 5 in control and AOM-injected, respectively), the effects being separated by a return to control level at 12 h and 18 h post-AOM (Fig. 3B). The level of AQP4 M1 isoform mRNA decreased by 47% (from 1.03 ± 0.28% in control to 0.56 ± 0.19) at 24h post-AOM (n = 7 and 3 in control and AOM-injected, respectively), whereas AQP4 M23 isoform mRNA dropped by 96% (from 1.035 ± 0.3 in control, to 0.04 ± 0.02) at 12h (n = 7 and 5 in control and AOM-injected, respectively), rose back to the control level at 18h (n = 7 and 3 in control and AOM-injected, respectively), and decreased again by 57% (to 0.44 ± 0.05) at 24h (n = 7 and 3 in control and AOM-injected, respectively) (Fig.3A). Protein level of PFN1 at 24 h post-AOM decreased by 36% (from 103.8% ± 16.95 in control to 66.75% ± 8.64; n = 8 and 7 in control and AOM, respectively) while it was not changed at the earlier time points (Fig.3C). The total protein level of ezrin remained unchanged throughout (Fig.3D). However, the analysis of the active, phosphorylated ezrin, imaged in Z-stacks from GFAP-labelled astrocytes, revealed its decrease by 27.4% in 24 h post-AOM (from 73.1 ± 27.2 to 53.3 ± 24.6; 40 astrocytes for control and 53 for HE, n = 4), with no changes at the earlier post-AOM times (Fig. 3E-F).

**Figure 3.**
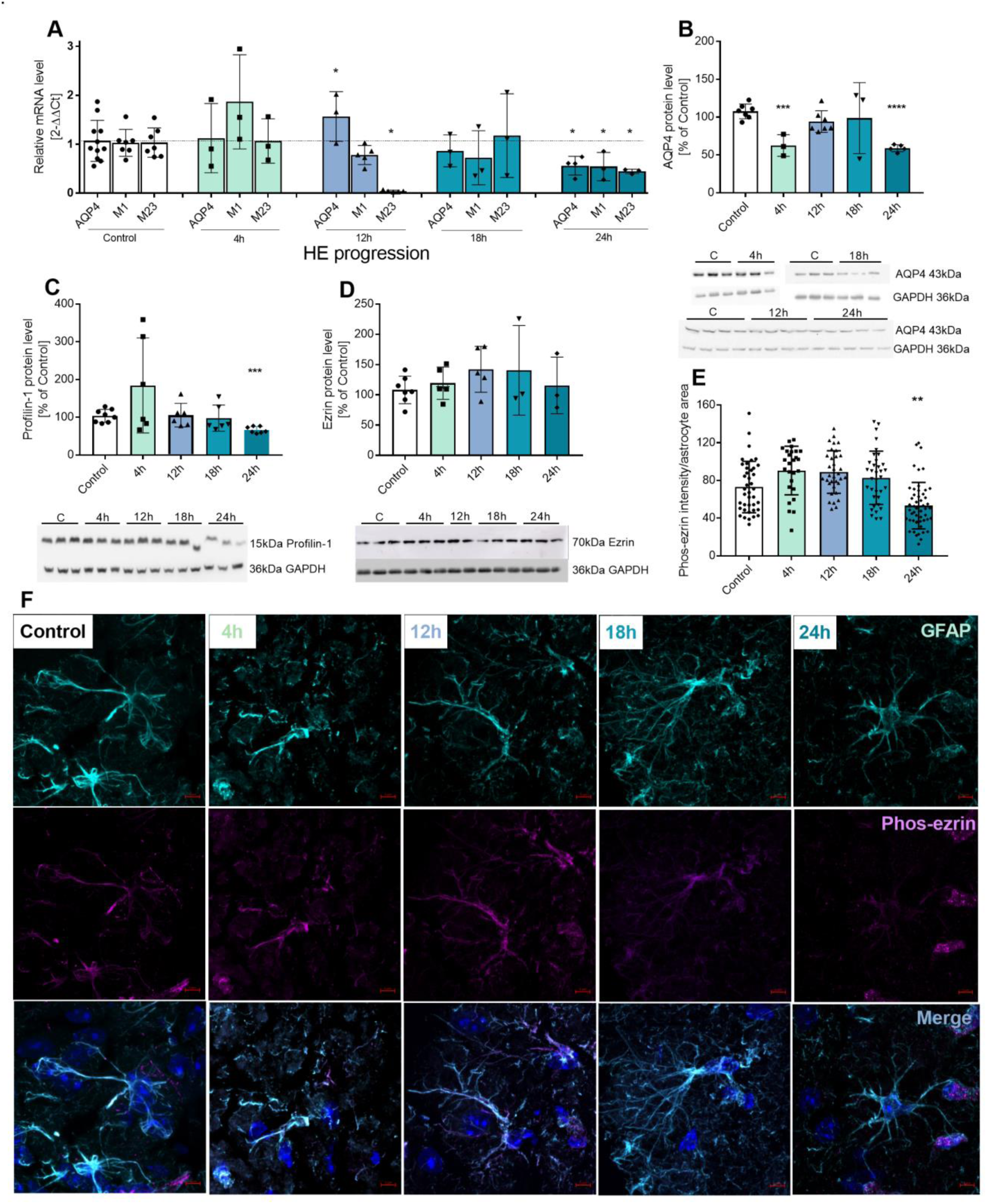
Changes of astrocytic aquaporin 4 (AQP4), profilin 1 (PFN1), ezrin, and phosphorylated-ezrin (Phos-ezrin) in the cortex of mice during HE progression. A: mRNA level of AQP4 and its M1 and M23 isoforms at 4, 12, 18, and 24 hours after AOM injection. N = 4-11, * p<0.05 (Mann-Whitney). B-D: Protein level of AQP4 (B), PFN1 (C), and ezrin (D) at 4, 12, 18, and 24 hours after AOM injection n= 3-7, * p<0.05, **p<0.0025, *** p<0.0003 (Mann-Whitney). E-F: Phos-ezrin (E – quantification of staining intensity, F – confocal microscopy images) in mice cerebral cortex at 4, 12, 18, and 24 hours after AOM injection.

### Remodelling of astrocytic arborisation in HE

The fluorescence intensity of TdT+ astrocytic processes was measured in 5 μm ring-shaped ROIs starting from the soma. Quantification of the ROI fluorescent intensity of maximum intensity Z-projected images (7 slices, 74 astrocytes, n = 3 mice for control group; 11 slices, 96 astrocytes, n = 4 mice for HE group) revealed increased intensity in two areas nearest to cell soma (0 – 5 μm and 5 - 10 μm), by 24.4% (from 31.1 ± 2.9 to 38.7 ± 1.9) and 17.3% (from 9.6 ± 1.0 to 11.3 ± 0.6), respectively. No changes were detected in 10 – 15 µm area while intensity was decreased in the areas further from the soma (15 – 20 μm, 20 – 25 μm, and 25 −30 μm), by 20.4% (from 4.7 ± 0.5 to 3.7 ± 0.4), 31.9% (from 3.6 ± 0.5 to 2.5 ± 0.5) and 37.5% (from 2.9 ± 0.5 to 1.8 ± 0.5), respectively (Fig. 4C). This pattern indicates hypertrophy of soma and primary branches and atrophy of distal astrocytic processes in HE mice.

**Figure 4.**
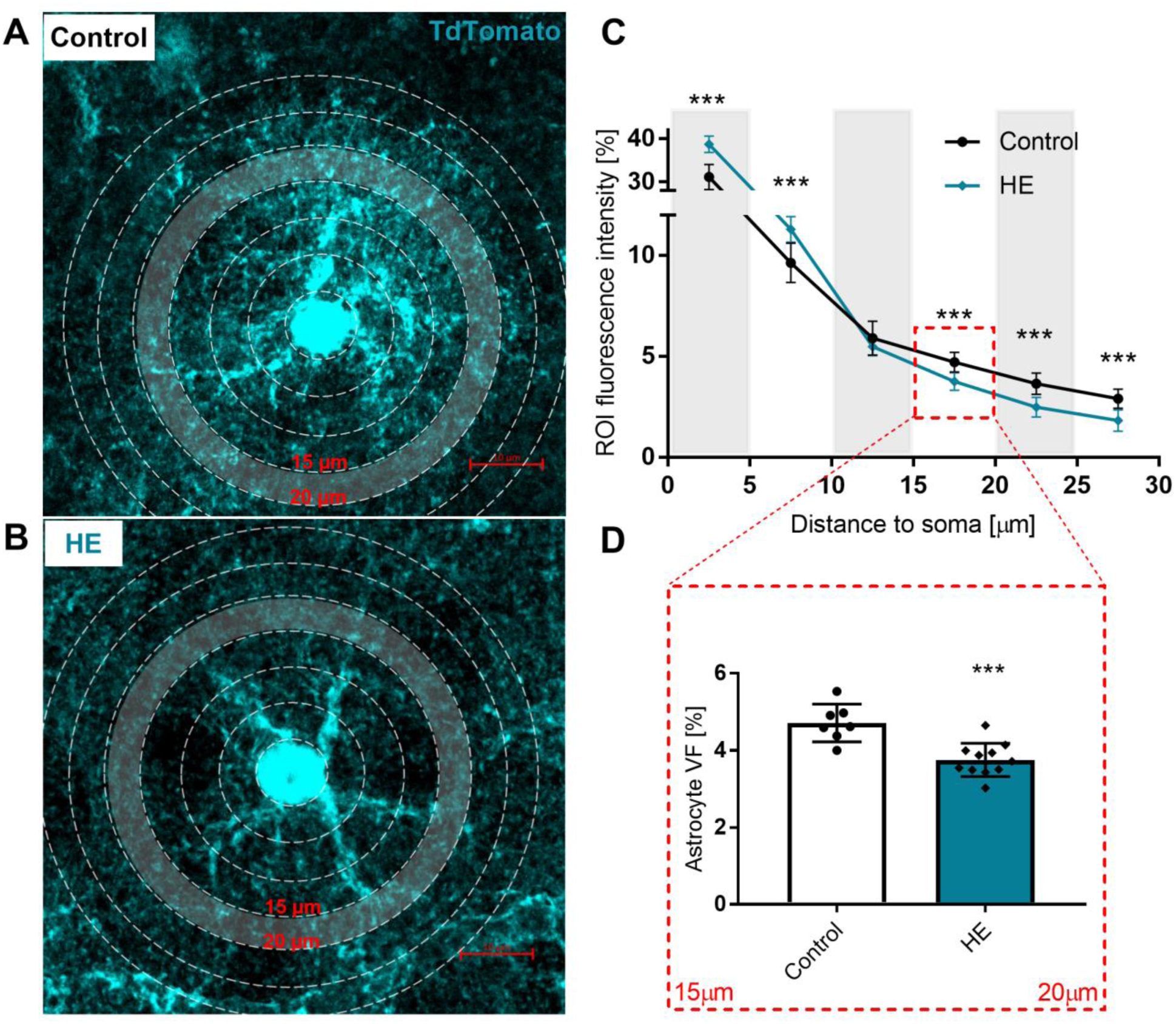
Immunofluorescent analysis of cortical astrocytic processes distribution in HE mice. A, B: Positioning of ROIs. C: Intensity of TdT fluorescence in 5 µm rings starting from the soma. D: Volume fraction of the area between 15 – 20 μm from the soma, indicating the fractional volume of the astrocyte. n=4 ***p<0.001 ANOVA with Dunnett’s post hoc test.

Volume fraction (VF) of leaflets, comprising fluorescence intensity from the entire area within 15 - 20 μm from the soma was analysed in Z-Stack images. Mean VF was significantly reduced in the HE mice (4.71 ± 0.48%, in control; 3.75 ± 0.43% in HE; p < 0.001, 7 slices, 74 astrocytes, n = 3 mice for control group; 11 slices, 96 astrocytes, n = 4 mice for HE group; two-sample t-test; Fig. 4D), indicating atrophy of astrocytic leaflets.

Sholl analysis was performed on traced skeletonised TdT+ astrocytes, with 2 μm intershell radii. Plot analysis (9 slices, 23 astrocytes, n = 3 mice from control group; 11 slices, 23 astrocytes, n = 3 mice from HE group) revealed decreased number of intersections between 10 μm to 36 μm from the soma in HE astrocytes; the highest number of intersections for control group was at 16 μm from the soma (19 ± 5 for control and 15 ± 4 for HE) while for HE group the highest number of intersection was detected at 14 μm from soma (18 ± 4 for control and 15 ± 4 for HE) (Fig. 5C). The length of the longest branch did not change (49.2 ± 5.6 for control and 46.2 ± 7.0 µm for HE) (Fig. 5D). Average number of intersections was significantly reduced in HE group (61 ± 13 for control and 34 ± 9 for HE) (Fig. 5E). 3D reconstructions from Z-stack also revealed substantial decrease in the complexity of arborisation in astrocytes from HE animals (Fig. 5F).

**Figure 5.**
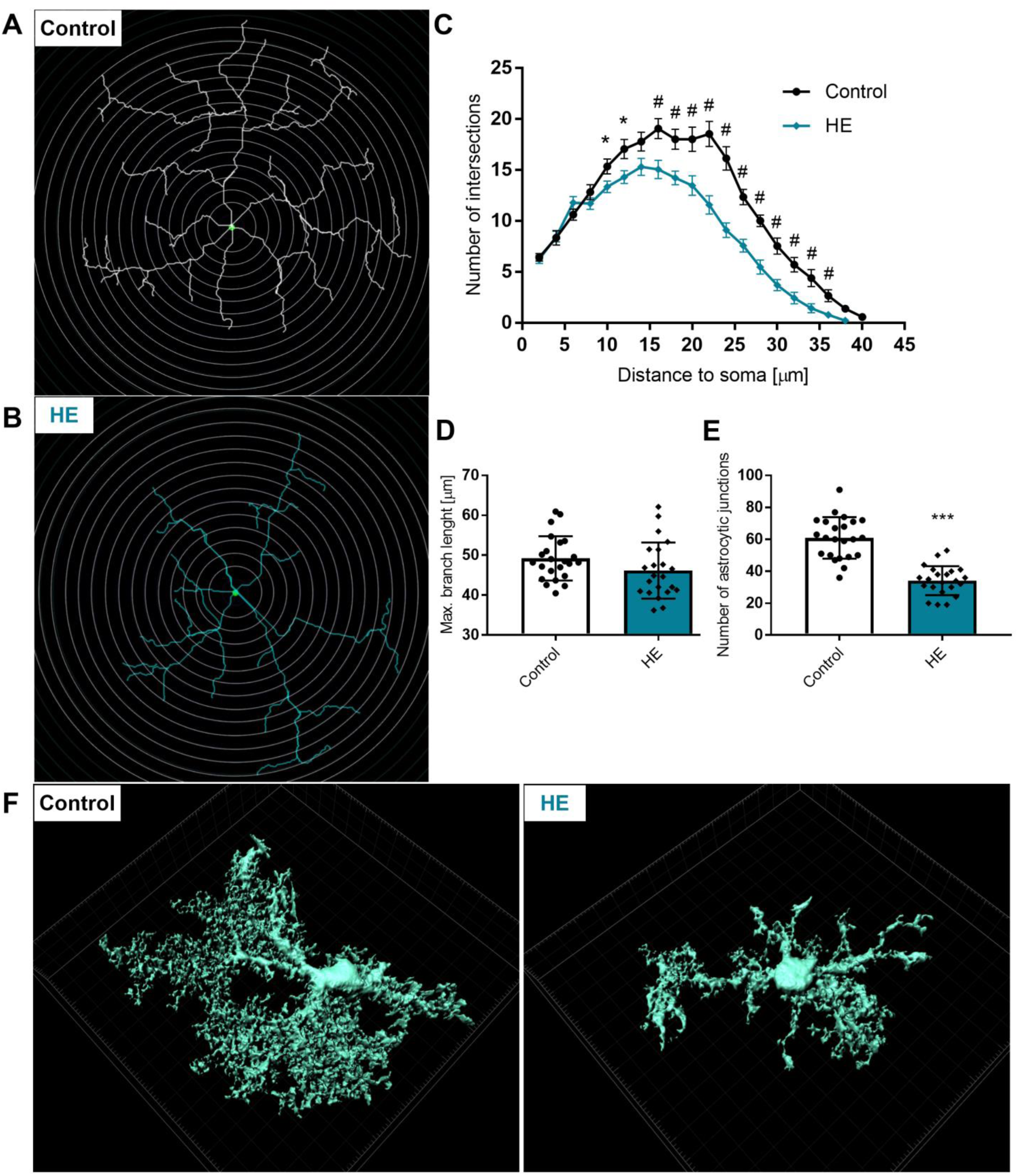
Sholl analysis of astrocytes. A-B: Sholl plot with 2 μm interval of control and HE astrocyte C: Sholl plot of C and HE astrocytes, D: longest branch length from soma centre, E: number of astrocytic junctions, F: Representative 3D rendered reconstruction of control and HE astrocytes.

### Rearrangement of astrocyte-neuronal synaptic landscape in HE

The volume of the neurone sharing the space occupied by the astrocytic branch was quantified by analysing Z-stacks TdT+ astrocyte branches and MAP2-immunostained neurons with a 3D surface rendering. Morphometric changes in the astrocyte-neuronal landscape revealed the global rearrangement of the morphological relations between astrocytic processes and neurones, with significant changes in astrocyte-neurone volume ratio of shared space. Space occupied by a single representative astrocytic arbour (including branches and leaflets) was analysed. We quantified astrocyte overlap with neighbouring neurones (MAP2-immunostaining) in control (72 analysed branches, 8 slices, n = 3 mice) and HE mice (99 analysed branches, 11 slices, n = 4 mice) (Fig. 6A, B). We observed a significant decrease in the astrocytic volume of individual branches from 2918 ± 718 μm^3^ in control to 1861 ± 378 μm^3^ in HE mice (Fig. 6D), which was accompanied by a decrease in astrocyte-neurone volume ratio in the shared space by ∼28% (1.56 ± 0.4 for control and 1.12 ± 0.16 for HE) (Fig. 6C).

**Figure 6.**
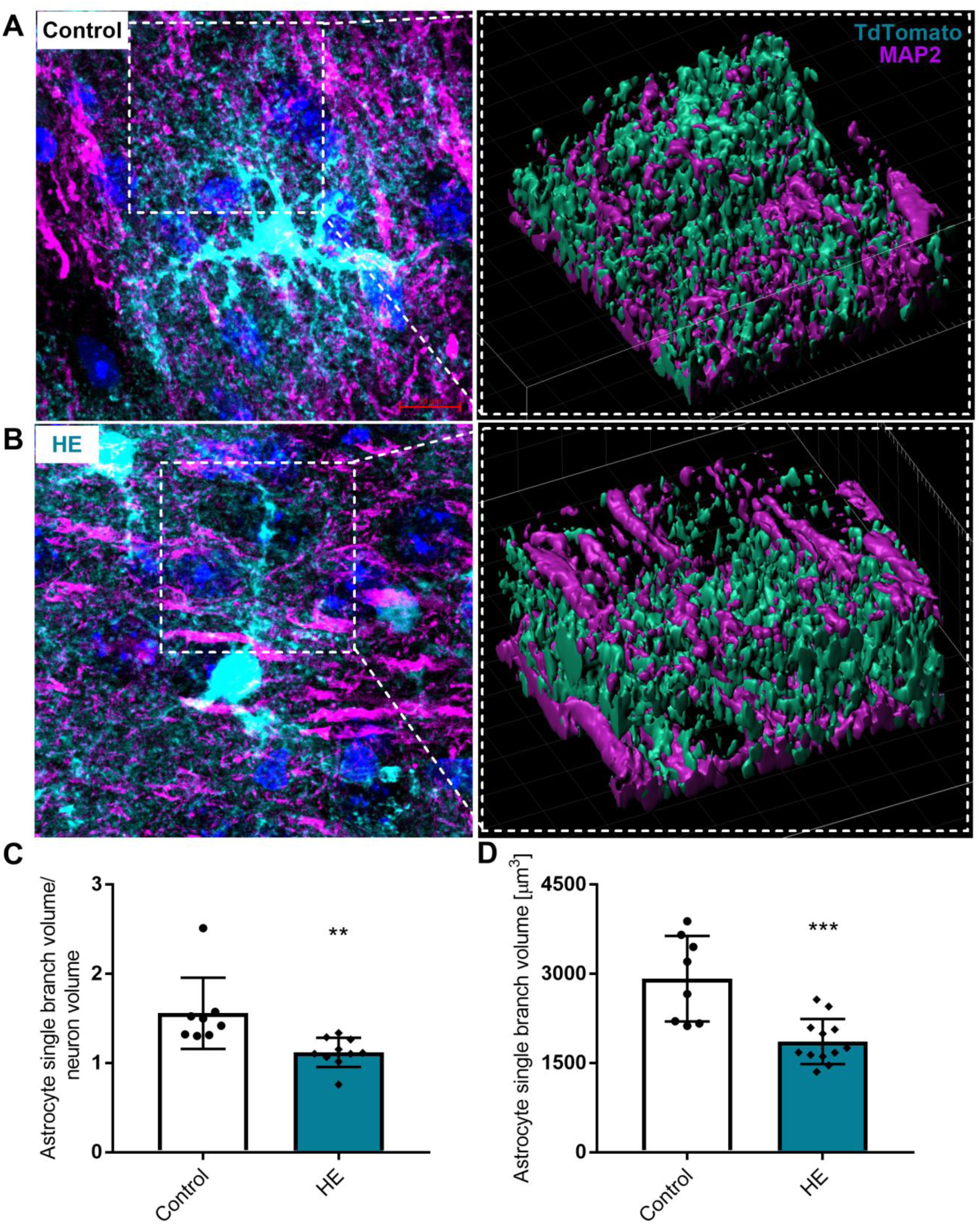
Rearrangement of astrocyte-neuronal landscape in HE. 3D reconstruction of astrocytic branch (TdT+) and neurones (MAP2+) from A: control and B: HE mice. C: Average volume ratio of the individual astrocyte branch to neurons. D: astrocytic individual branch volume, n=4, 80 sections from the group, ** p<0,01, ***p<0.001 ANOVA with Dunnett’s post hoc test.

### Neuronal decline in acute HE

#### EEG alterations in HE

Cortical EEG recording revealed decreased total EEG power, starting from 12 h post-AOM administration (Fig. 7A), with substantial decrease from 100% at baseline to 59% ± 22.62, 56.81% ± 30.45 and 49.71% ± 23.7 in HE mice (n = 5) at 12, 18, 24 h post AOM, respectively. A similar pattern of brain activity decline was observed when the EEG spectra were split into separate bands – a decrease of ʘ, α, and β waves became apparent from 12 h post-AOM, whereas γ and, HFO waves decreased starting from 18 h post-AOM (Fig. 7A).

**Figure 7.**
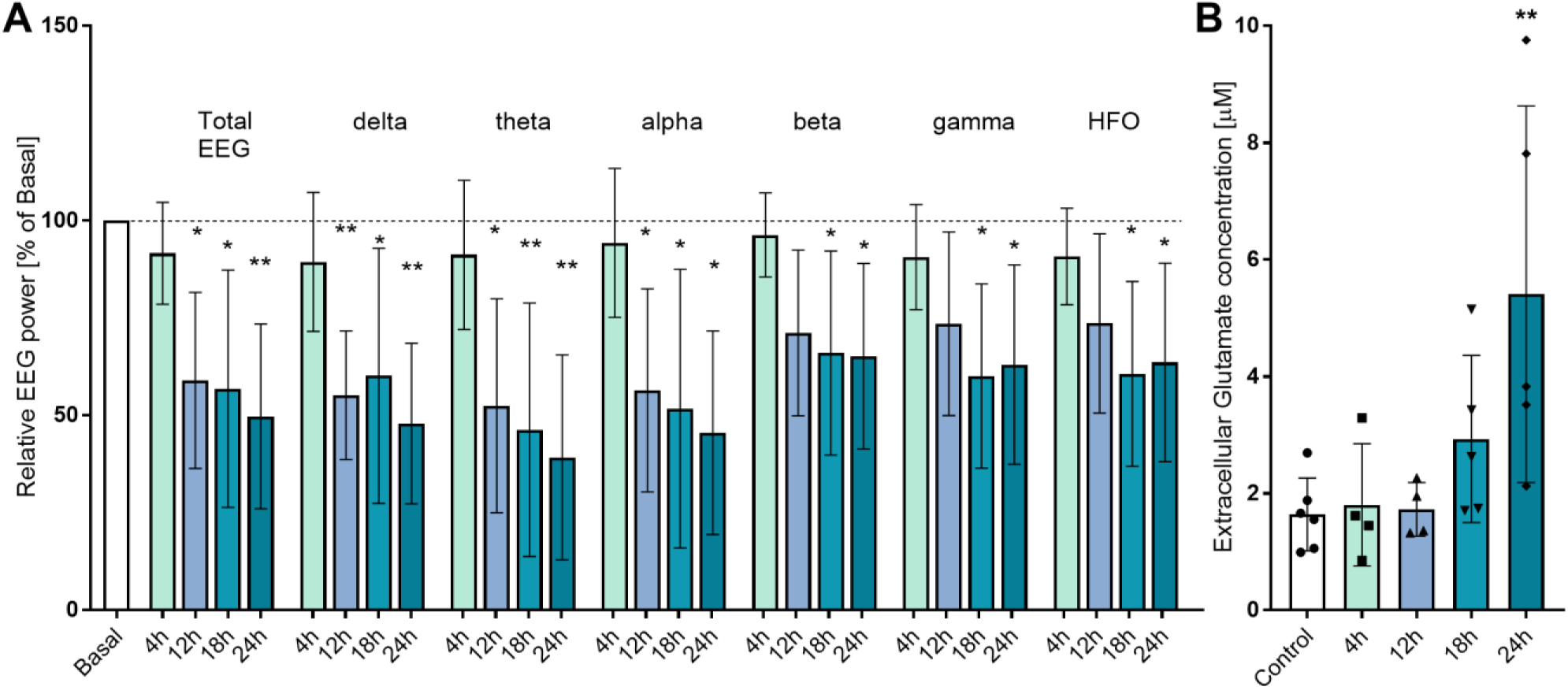
Progressive decline in neural functions in ALF mice A: The relative EEG power in mice cortex at 4, 12, 18, and 24 h after AOM injection. B: Extracellular glutamate concentration in microdialysates from the frontal cortex of mice 4, 12, 18, 24 hours after AOM injection, n=4-5, **p<0.01 ANOVA with Dunnett’s post hoc test.

#### Extracellular glutamate level in HE brain

From about 18 h after AOM administration, glutamate levels showed a tendency to increase (without statistical significance), which change persists later, in HE mice we observed substantial increase from 1.64 ± 0.62 μM in control (n = 6) to 5.41 ± 2.44 μM in HE mice (n = 5) (Fig. 7B).

## Discussion

Morphological changes of astrocytes associated with toxic encephalopathies were discovered in 1912 by Carl Von Hösslin and Alois Alzheimer (von Hosslin & Alzheimer, 1912) who analysed postmortem tissues of a patient with Westphal-Strümpell syndrome, a toxic copper encephalopathy, known today as Wilson’s disease. Ammonium toxicity as a factor underlying HE was described in early 20s century, after analysing the meat intoxication in Eck fistula dogs (Sherwood et al., 1916). Association between ammonium accumulation in the brain and astrocytic remodelling was first established about 50 years later (Norenberg & Lapham, 1974), whereas detailed morpho-functional analysis of the astrocytic changes remained incomplete until present.

In this study, we employed the AOM model of ALF, accepted in acute HE studies, as it sufficiently reproduces neurological and neurobehavioural impairments (DeMorrow et al., 2021; Popek et al., 2018). To characterise the disease progression at defined time points, we broadened the morphological and biochemical characteristics of the model and confirmed gradual liver failure after AOM injection.

Here we demonstrate, to our knowledge for the first time, morphological remodelling of astrocytes including retraction of astrocytic leaflets from synapses in the cerebral cortex of mice with HE. We also quantified changes in ezrin, PRF1, and AQP4 associated with morphological plasticity (Ciappelloni et al., 2019; Lavialle et al., 2011b; Molotkov et al., 2013; Schweinhuber et al., 2015). Most significantly, in the HE, a substantial decline in AQP4, PRF1, and phosphorylated ezrin, correlated with (and arguably instigated) the atrophy of astrocytic perisynaptic leaflets suggesting a causal nexus between these phenomena. We provide evidence that the astrocytic remodelling may be linked to dyshomeostasis of neurotransmission, reflected by changes in EEG, and by a substantial increase in extracellular glutamate.

Astrocytic malfunction in the HE was mainly thought to be linked to brain oedema, which often represents the primary cause of death in this disease (Scott et al., 2013). The role of astropathology is, however, wider, as loss of astrocytic homeostatic support and impaired ability of astrocytes to control brain ions, energy, and neurotransmitter homeostasis contributes to synaptic malfunction, neuronal damage, and ultimately to psychiatric and cognitive clinical presentations (Verkhratsky et al., 2023). Our previous studies established aberrant neurotransmission associated with abnormal calcium buffering in presynaptic terminals (Popek et al., 2018; Hamdani et al., 2021). The present study highlights, to our knowledge for the first time, the concurrence of astrocytic structural remodelling and synaptic malfunction in the progression of HE.

Astrocytes actively control neuronal function by instructing synapse formation, plasticity, and remodelling (Allen & Eroglu, 2017; Baldwin & Eroglu, 2017; Verkhratsky & Nedergaard 2018). Their essential functions at the synapse are intimately linked to complex astrocytic morphology (Oberheim et al., 2006). From the above perspective, the term primary astrogliopathy, emphasises the key role of astrocytes in the development of HE, similarly to other diseases with a malfunctional astrocyte component (Verkhratsky & Butt, 2023). Alzheimer type II astrocytes, although being idiosyncratic to HE, represent only a minor subpopulation of astroglia in the normal and pathological brain.

Since the ramified morphology of astrocytes determines their close structural association with synapses (Semyanov and Verkhratsky, 2021), astrocytic morphological plasticity (physiological or pathological) influences synaptic transmission (Nagai et al., 2021; Santello et al., 2019). Morpho-functional aberrations of astrocytes were described in neurodegenerative, neuropsychiatric, and neuromuscular diseases characterised by slow progression (Verkhratsky & Butt, 2023; Verkhratsky et al., 2023). However, in rapidly developing pathologies that primarily affect astrocytes, including HE, neither the details of the remodelling process, nor association with astrocyte dynamics at the synaptic sites of astrocyte-neurone interaction have been hitherto elucidated. In our study, by in depth quantification of astrocyte morphology, we assess the HE cortical landscape from the astrocyte processes remodelling perspective. First, we documented a significant increase in the total astrocyte area of the HE frontal cortex neuropil sections. Ultrastructural analysis agrees well with our previous reports of an increase in the astrocytic area in the HE brain, reflecting changes in astrocyte morphology associated with swelling (Hamdani et al., 2021). We further quantified the areas occupied by perivascular astrocytes vs astrocyte branches localised at a distance from the vessels (brain neuropil). Irrespective of whether we quantify astrocytic endfeet *in toto* or their branches and leaflets, HE is linked with a significant increase in the territory occupied by astrocytes.

Further, to address astrocyte complexity, we performed an immunofluorescence analysis of the distribution of astrocytic processes. The comparison of the astrocyte leaflet volume fraction (VF), revealed noticeable decrease in the VF starting from around 15 μm distance from soma in HE mice. At the same time, the VF was higher closer to the soma.

Astrocytic structural remodelling occurs in response to different stimuli (e.g., osmotic stimulation, stress, neuronal activity, etc.). Changes in plasticity-related proteins AQP4, the phosphorylated protein ezrin, and PFN1, correlate with astrocyte morphology changes. Astrocyte sensitivity to the osmotic changes results from their ability to rapidly take up excessive water and ions through AQP4 and associated ionic channels. Indeed, the involvement of AQP4 in the brain volume changes in HE was often (Tang & Yang, 2016), albeit inconsistently (Saadoun & Papadopoulos, 2010) reported. We found a decrease in AQP4 protein, phosphorylated ezrin, and PRF1 in HE brains. Both PRF1 and ezrin are required for the morphological plasticity of leaflets (Derouiche & Frotscher, 200; Badia-Soteras et al., 2023). Translocation of AQP4 within the astrocyte membrane near adjacent neurones may indirectly affect extracellular levels of neurotransmitters by changes in extracellular volume during synaptic activity. In line with the above, the variations in the expression of mRNA coding for the M23 isoform may reflect, or contribute to HE-induced astrocytic adaptive change in astrocyte morphology. Leaflet atrophy, observed in this study, translates into reduced synaptic coverage, which potentially may affect neurotransmission. Quantification of astrocytic leaflets surrounding the perimeter of cortical synapses demonstrated a lower number of synapses in close contact with a single astrocytic leaflet, in favour of contact with two leaflets. Further, distances between the synaptic active zone and the closest astrocytic membrane increased by ∼25% in HE. Increase in the number of leaflets contacting synapses may be a compensatory response to aberrant neuronal activity aimed at more effective astrocyte-neurone communication or is a consequence of astrocyte failure in response to swelling.

Reduced EEG activity is a consistent feature of advanced, symptomatic HE in human patients (Sutter et al., 2013) and experimental animals (Cittolin-Santos et al., 2019; Rangroo-Thrane et al., 2012; van der Rijt et al., 1990), reflecting decreased synaptic transmission. Previous studies highlighted the inefficient recruitment and/or impaired trafficking of synaptic vesicles to synaptic active zone as one of the possible molecular mechanisms underlying decreased neurotransmission in the HE (Potvin et al., 1984; Popek et al., 2018.

Increase of extracellular glutamate concentration measured in the brain microdialysates of HE rats is consistent with findings in human HE patients (Tofteng & Larsen, 2002) and different rodent HE models (Cittolin-Santos et al., 2019; Vogels et al., 1997; Cauli et al., 2008). Reduction of EEG power with increased extracellular concentration of glutamate has been reported earlier in other animal HE models (Bosman et al., 1990; Cittolin-Santos et al., 2019).

Glutamate spillover frequently observed in HE is related to the deficiency of astrocytic glutamate uptake observed in hyperammonemic conditions (Dabrowska et al., 2018). Accumulation of extracellular glutamate due to astrocytic homeostatic failure impairs neurotransmission, affects neuronal excitability, and instigates excitotoxicity.

Astrocytic changes in HE brain are complex; in addition to an overall increase in the volume of astrocytes at the level of soma and primary branches, most likely reflecting astrocyte oedema, we found decreased complexity of astrocytic arborisation manifested by atrophy of distal branches and leaflets. These morphological changes, which are remarkably rapid (to compare, reactive astrogliosis in response to brain trauma is initiated 24 – 36 hours after the insult (Verkhratsky et al., 2023), are associated with reduced synaptic coverage and arguably loss of astrocytic homeostatic support.

## Conclusions

In the present study, we demonstrated that cortical astrocytes in the HE induced by ALF undergo complex morphological alterations. The soma and primary branches swell, whereas complexity of distal arborisation is reduced, mainly by rapid atrophy of distal branches and leaflets. These results in the reduced astrocyte synaptic coverage that translates into decreased neuronal activity. Thus rapid emergence of aberrant astrocytes presents the fundamental element in the pathogenesis of the HE.

## Acknowledgments

We want to thank Dr. Marek Pawlik for his help in EEG recordings.

The authors express special gratitude to Prof. Maciej Garstka and Dr. Radoslaw Mazur for help with cell morphology analyses.

The fluorescence intensity of Phos-ezrin was performed in Laboratory of Advanced Microscopy Techniques, Mossakowski Medical Research Centre, Polish Academy of Sciences.

## Funding

Financial support was provided by statutory funds No. 13 from Mossakowski Medical Research Institute, Polish Academy of Sciences, Warsaw, Poland (MP, MO-P, JA, MZ), jointly managed with the National Science Centre in Poland within MINIATURA 3 (no 2019/03/X/NZ4/00623; MO-M).

## Author contributions

Conceptualisation: MZ, MO-M, MP; Methodology: MP, MO-M, ŁS, MK, MZ; Investigation: MP, MO-M, MK; Visualisation: MP; Funding acquisition: MZ, MO-M; Supervision: MZ; Writing – original draft: MZ, MO-M, MP; Writing – review & editing: MZ, JA, AV.

## Competing interests

Authors declare that they have no competing interests.

## Data and materials availability

Data are available upon reasonable request.

## Notes

### Competing Interest Statement

The authors have declared no competing interest.

